# Toward One-Shot Learning in Neuroscience-Inspired Deep Spiking Neural Networks

**DOI:** 10.1101/829556

**Authors:** Faramarz Faghihi, Hossein Molhem, Ahmed A. Moustafa

## Abstract

Conventional deep neural networks capture essential information processing stages in perception. Deep neural networks often require very large volume of training examples, whereas children can learn concepts such as hand-written digits with few examples. The goal of this project is to develop a deep spiking neural network that can learn from few training trials. Using known neuronal mechanisms, a spiking neural network model is developed and trained to recognize hand-written digits with presenting one to four training examples for each digit taken from the MNIST database. The model detects and learns geometric features of the images from MNIST database. In this work, a novel biological back-propagation based learning rule is developed and used to a train the network to detect basic features of different digits. For this purpose, randomly initialized synaptic weights between the layers are being updated. By using a neuroscience inspired mechanism named ‘synaptic pruning’ and a predefined threshold, some of the synapses through the training are deleted. Hence, information channels are constructed that are highly specific for each digit as matrix of synaptic connections between two layers of spiking neural networks. These connection matrixes named ‘information channels’ are used in the test phase to assign a digit class to each test image. As similar to humans’ abilities to learn from small training trials, the developed spiking neural network needs a very small dataset for training, compared to conventional deep learning methods checked on MNIST dataset.

## Introduction

The human brain has demonstrated amazing cognitive capabilities to learn and recognize complex visual patterns in noisy context [1, 2]. Information processing in the human brain is done via activation of sensory neurons and subsequently sending the inputs into cortical neurons that leads to complicated spiking pattern of large neuronal population to make decision or store the information [3]. Neurons in the cortex are sparsely connected via dynamical synapses that are weakened or strengthened [4, 5] by some mechanisms, such as activity-dependent or retrograde signaling from other neurons for communication of different regions of the brain. Understanding such network and molecular mechanisms and then implementing them in future artificial systems may lead to brain-like machines to perform complex tasks [6, 7]. One of these mechanisms is back-propagation of action potentials that occurs in some neurons in which after generating an action potential by a neuron down the axon, another electric pulse is generated from the soma and propagates toward to dendrites. These signals seem to carry information to other neurons about the overall state of corresponding neurons [8]. Activity-induced retrograde messengers as simple molecules or short peptides are crucial for the formation of some synapses in some regions of the brain through learning and memory [9]. The role of back propagation of action potentials as principal mechanism of triggering diffusion of retrograde signals and their role in network organization has been shown [10]. However, the learning processes in the brain do not appear to use back propagation of error signals as it uses a deep neural network architecture. Deep neural networks are artificial neural networks that are composed of more than two neural layers.

During neural development, synapses are over-produced and then eliminated over time. This neuronal event is known as ‘synaptic pruning’. Developmental synapse formation as an activity-dependent process play an important role in healthy brain where synaptic pruning may be mostly regulated by activity-dependent mechanisms [11, 12]. However, for engineered network design, connections are usually added over time from an initially sparse topology. To design engineered networks, adding connections that will soon be removed is considered wasteful [13]. Artificial intelligence society has been considering neuroscience as a rich source of inspiration for developing cognitive systems doing complex tasks that currently artificial systems are not able to perform [14].

Models of cortical structure and function have inspired developing deep learning models but there are no remarkable similarity between biological and artificial deep networks [15, 16]. On the other hand, machine learning models can give explanations and assumptions for how the brain may achieve complicated tasks in uncertain environments [17]. However, the main challenge to achieve these goals is the nature of different fundamental algorithms of machine learning (that use matrixes of data) and the mechanisms underlying cognitive capabilities of the brains.

Deep learning, which is a field of machine learning, allows researchers in different field to process information of interests by sending input information to multiple artificial neural layers which can be trained by large set of data to recognize features of complex inputs (voice, image, etc.) [15]. In recent years, very important progresses have been accomplished in neuroscience-inspired algorithms [18]. Spiking Neural Networks (SNNs) have demonstrated capabilities for information processing of data of different sources [19]. In SNNs, sparse and asynchronous binary signals are used as communications tool and synaptic weight between neurons is subject to time and activity dependent parameters. SNNs have successfully been implemented to model complicated neuronal dynamics underlying cognition [20]. Remarkable success of deep learning in machine learning for analysis of data from different sources has motivated computer scientists to view deep SNNs as a more efficient replacement of conventional DNNs [21]. A challenge to apply SNNs in deep neural networks is the discontinuous nature of neural communication in SNNs as generating spikes over time. Therefore, direct application of back-propagation algorithm in SNNs as used in conventional neural networks is an important challenge for the future spiking deep learning methods [21]. For a review on the challenges and applications of SNNs in deep neural networks, we refer to [21]. One of the attempts of modern deep learning research is to develop spiking deep networks by understanding how networks of biological neurons learn to achieve visual tasks and perform feature extraction [22].

One of the interesting example of human capabilities to recognize noisy and complicated information is the hand-written digits recognition that is currently considered as an evaluation benchmark for deep learning methods. Visual system of human extracts features from noisy and incomplete information in images of the dataset. Handwritten digit recognition is critical in some machine learning application, such as postal mail sorting and bank check processing. The complexity of the problem arises from the fact that handwritten digit images are not always of the same size, width and orientation, so the general problem would be recognizing the similarity between images of the digit classes [23].

MNIST dataset is a database for evaluating and improving machine learning models on the handwritten digit classification problem. MNIST is widely used as an excellent dataset for evaluating models as well as deep learning and machine learning algorithms [24]. For this purpose, several machine learning algorithms have been used and evaluated their efficiency on MNIST database, including multi-layer perceptron, SVM (support vector machines), Bayes nets and Random forests [25]. More specifically, different artificial neural network architectures have used MNIST database for digit recognition task, while the best performance on classification of the test dataset has been done by deep neural networks [26]. Deep SNNs (DSNNs) are brain-inspired information processing systems, which have shown interesting capabilities such as fast inference and event-driven information processing which make them excellent approaches for deep neural network architectures [27]. Event-driven means that SNNs generate spikes in response to stimulation from other neurons and show very small firing activity when they receive sparse inputs, such strategy results in power-efficient computing [28]. DSNNs have been developed for supervised learning [29], unsupervised learning [30, 28] and reinforcement learning paradigms [31].

In this work, we present a new approach that is inspired by synaptic and systems neuroscience for classification of hand-written digits. The motivation of the model is the human approach to distinguish digits by features selection (vertical, horizontal lines and circles that compose digits). The work present a new learning rule that is inspired by biological back-propagation and synaptic pruning that are both known in brains. The main goal of the model is to test a hypothesis on how the brain may learn digits shape with a few training examples and its application in developing future deep learning methods.

## 2. Method

One of the main aims of neuroscience-inspired machine learning is to develop neural networks that are capable of pattern recognition using small training sets. The human neocortex has demonstrated high performance for patterns recognition tasks by presenting a few training sets while artificial systems needs a very large training sets (e.g. deep learning techniques). One of well-known example of such tasks is the recognition of hand-written digits. Toward developing neuroscience-inspired machine learning, we need to gain a good level of mechanistic understanding of neuronal computations in processing units in the brain and in local networks as well. More specifically, synaptic mechanisms and how time dependent learning mechanisms are involved in building recognition tasks are essential to achieve goals. In this work, we have developed a Deep Spiking Neural Network (DSNN) that can demonstrate similar pattern recognition capability for hand-written digits recognition. In the human brain, the pattern of vertical, horizontal and circles and their combinations that construct digits is learned to assign a digit to a picture. The logic of learning digits by the brain seems to be fundamentally very different from learning algorithms used in deep learning methods. We have developed a new approach to use both simplified synaptic mechanisms and dynamics of activity-dependent synaptic connections to test on MNIST database by training with a small training set. The basic synaptic mechanism used in this work is back-propagation-based learning rule and synaptic pruning.

The network is composed of three layers (Figure 1). The input to the network is based on the MNIST dataset as images of digits 0 to 9. Handwritten digits are images in the form of 28*28 intensities matrixes. The SNN consists of neuronal layers that contain Integrate and Fire neurons (IF-neurons). At the first stage, information in the images are partitioned shown in Figure 2B. Each image as 28*28 pixels are partitioned into two parts by a hypothetical vertical or horizontal or orthogonal line to divide equally each image. Then each pixel of selected-set of pixels as horizontal, vertical or orthogonal segments are considered as ‘spike train generators’ with firing rates proportional to the intensity of the pixels (the first layer). Each selected set of pixels in vertical, horizontal and orthogonal lines are fully connected to a single IF-neuron for 1000 *ms*. For this purpose, the maximum pixel intensity of 250 is divided by 4 to generate firing rates between 0 and 63 HZ [30]. In addition, we did a preprocessing on the images before applying spiking based method as follows. We normalized the pixels by a simple rule as considering the pixels’ values equal or higher than 0.4 to 0.4 (Figure 2A). Consequently, the 224 neurons (28*8, 8 partitions and 28 set of neurons in the first layer) that receive the spikes from the first layer construct the second layer. The third layer composed of 150 neurons (arbitrarily number in this study) are fully connected to the second layer with random synaptic weights values between 0 and 1 before training phase where the weights are dynamically changes according to the learning rule.

**Figure 1.**
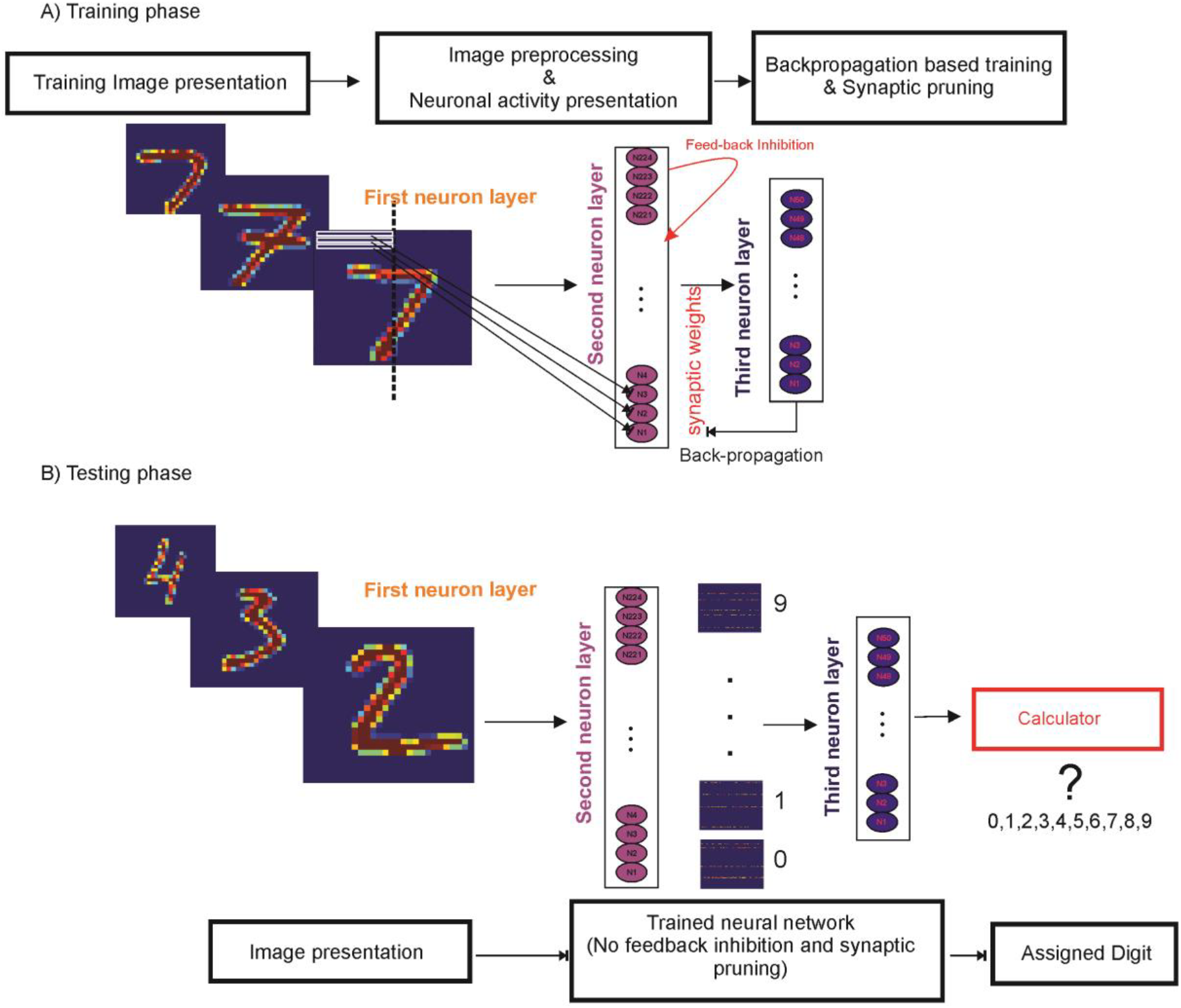
Schematic representation of the network architecture. **A.** Training phase. For each digit (0-9) a number of samples are selected from the dataset and information of each picture (28*28 pixels) is represented as neuronal activity of 224 ‘integrate and fire neurons’ (first layer). In each epoch of presenting the sample picture the spiking activity of the neuron in the first layer is sent to the second neuronal layer (150 integrate and fire neurons). Throughout training, the synaptic weights between two layers that are initially randomly selected are changed according to the biological back-propagation based learning rule defined in this work and eventually synaptic weights lower than the threshold are set to zero (to model synaptic pruning). The feedback inhibition in the second layer that controls the activity of the network and prevents over firing of the neurons. **B.** Testing phase. After training the network (12 epoch for each training image) for all digits, images are presented to the network to assign one digit to each picture and check whether it is correct. Each image is transformed into the activity of first neuronal layer and then 10 different connectivity matrixes are used to send information into the second layer. The input of the second layer is computed for different connectivity matrixes and then the maximum value of the inputs is used to assign a digit for each presented image.

**Figure 2.**
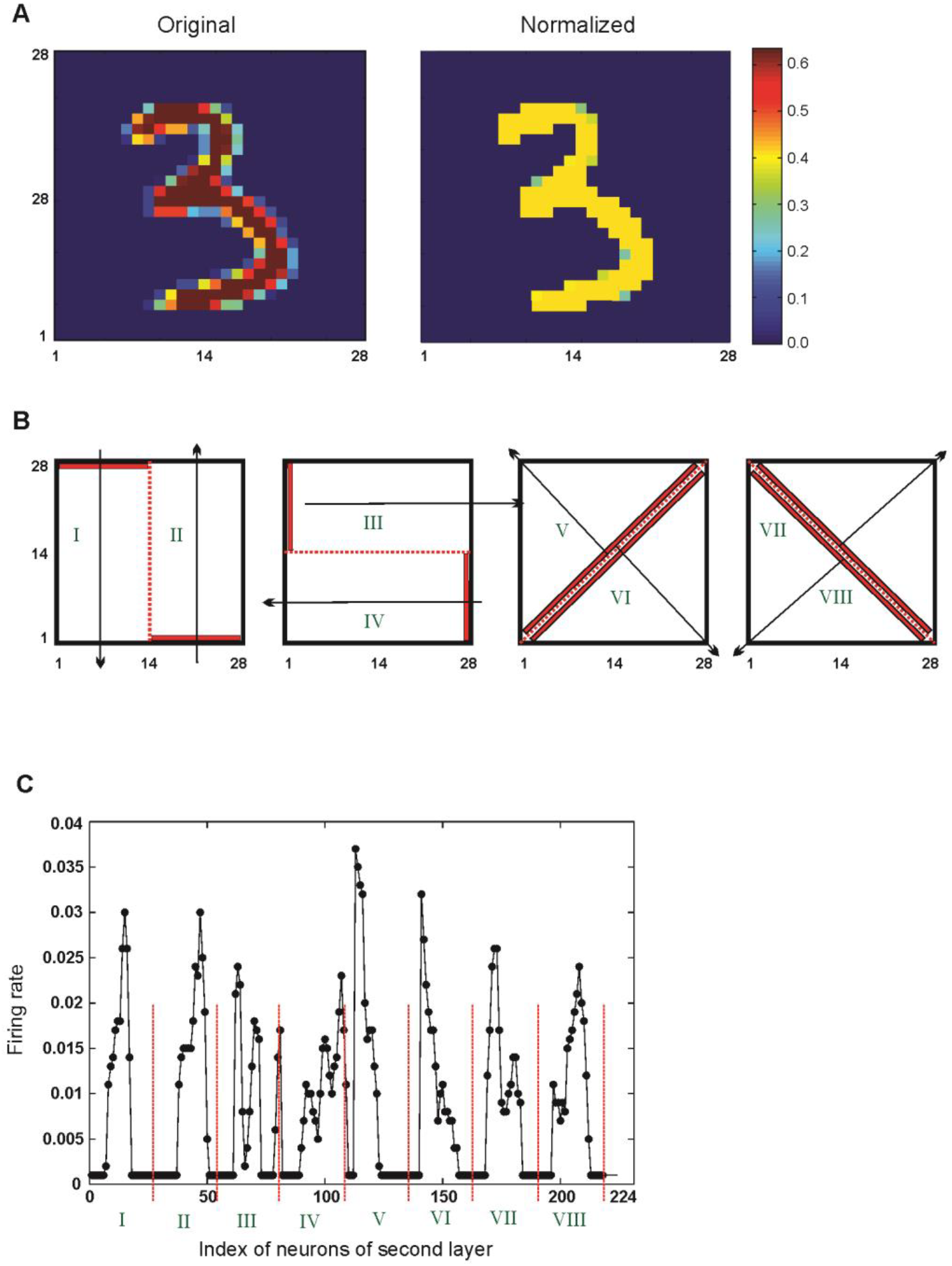
**A.** Image preprocessing. The hand-written images are presented as pixels value ranging from zero to 250 which is normalized between zero and 0.4. **B.** Representation of image’s information as neuronal activity of neuron in first layer. For this purpose, the image is divided into two regions as shown by vertical, horizontal and orthogonal lines of pixels. Each region is represented as the spiking activity of 28 neurons. **C.** Average firing rate of a neuron in the second layer for 1000 time bin image presentation.

Each neuron in the second and third layers shows spiking activity as IF-neurons (**Equation.1**).

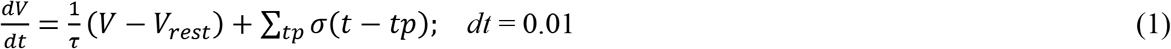

The spiking activities of each neuron in the second layer are controlled by an inhibitory neuron; its activity is described by **Equation.2**.

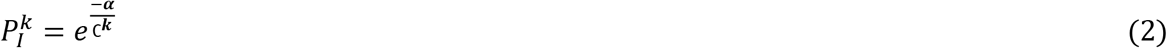

Where 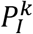 is the probability to inhibit spike generation by neurons in the second layer. *α* is the inhibition parameter value and ∁^*k*^ denotes the average activity of neuron *k* of the second layer.

During the training phase, after a full stimulation time (500 *ms*) of the network by image presentation, the average firing rate of total neurons in the third layer is measured. The difference of average firing rate of each pair of neurons in the second layer and the third layer is measured and shown by ∆_(*i,j*)_ (**Equation.3).** The change in the synaptic weight of the pairs of neurons (∆*w*_(*i,j*)_) is calculated as **Equation.4**.

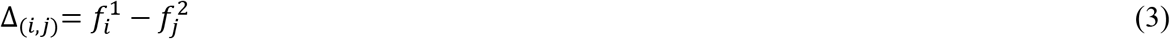

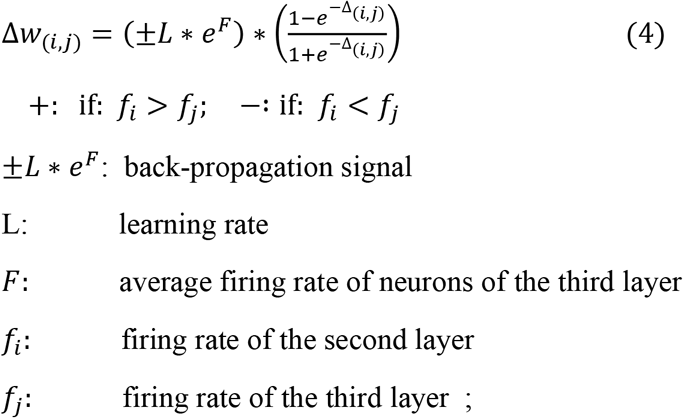

After presentation training images for each digit to the network, the test phase is done. The images from the MNIST dataset is presented to the train network where 10 different connectivity matrixes are used to measure the total input from the second layer into the third layer (**Equation.5**). The corresponding digit that shows maximum input into the third layer is considered as assigned digit (*k*) to the image (**Equation.6**).

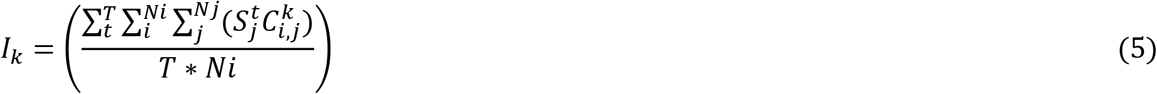

Where *T* is the epoch time, *Nj* is the number of neurons in the third layer, *Ni* is the number of neurons in the second layer, 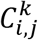 is synaptic weight matrix between the second and third layers and 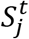 is the spiking of a neuron in the third layer.

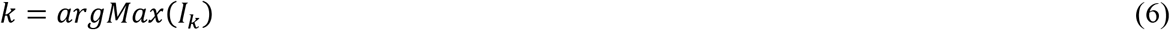

## Results

Each presented image to the network stimulates spiking activity of 224 neurons in the second layer. Average firing rate of these neurons for a sample of digit ‘3’ from the MNIST dataset are shown in Figure 2.C. Figure 2.C also shows the activity pattern of partitions defined in Figure 2.B.

One can expect that such observed patterns would be more or less very similar for the samples from the same digit class while shows different patterns of images for different digit classes. For this purpose, two images of digit ‘4’ and two images of digit ‘7’ were selected and the pattern of average spiking rate of the neurons in the second layer for pair of digits are shown in Figure 3.A, B. The spiking activity pattern of neurons in the second layer for images of different classes of digit are shown in Figure 3.C. The distance of the pattern as a measure of similarity of the patterns are shown in Figure 3.D. The results show that selected digits from the same class are more similar compared to images from different digit class. Regarding the variation in the hand-written digits in the dataset where digits are written with different intensities and thickness and the impact of image intensity on the spiking of neurons in the developed spiking network (Figure 4.A), feedback inhibition with different intensity value were used to control activity level of the neurons in the network.

**Figure 3.**
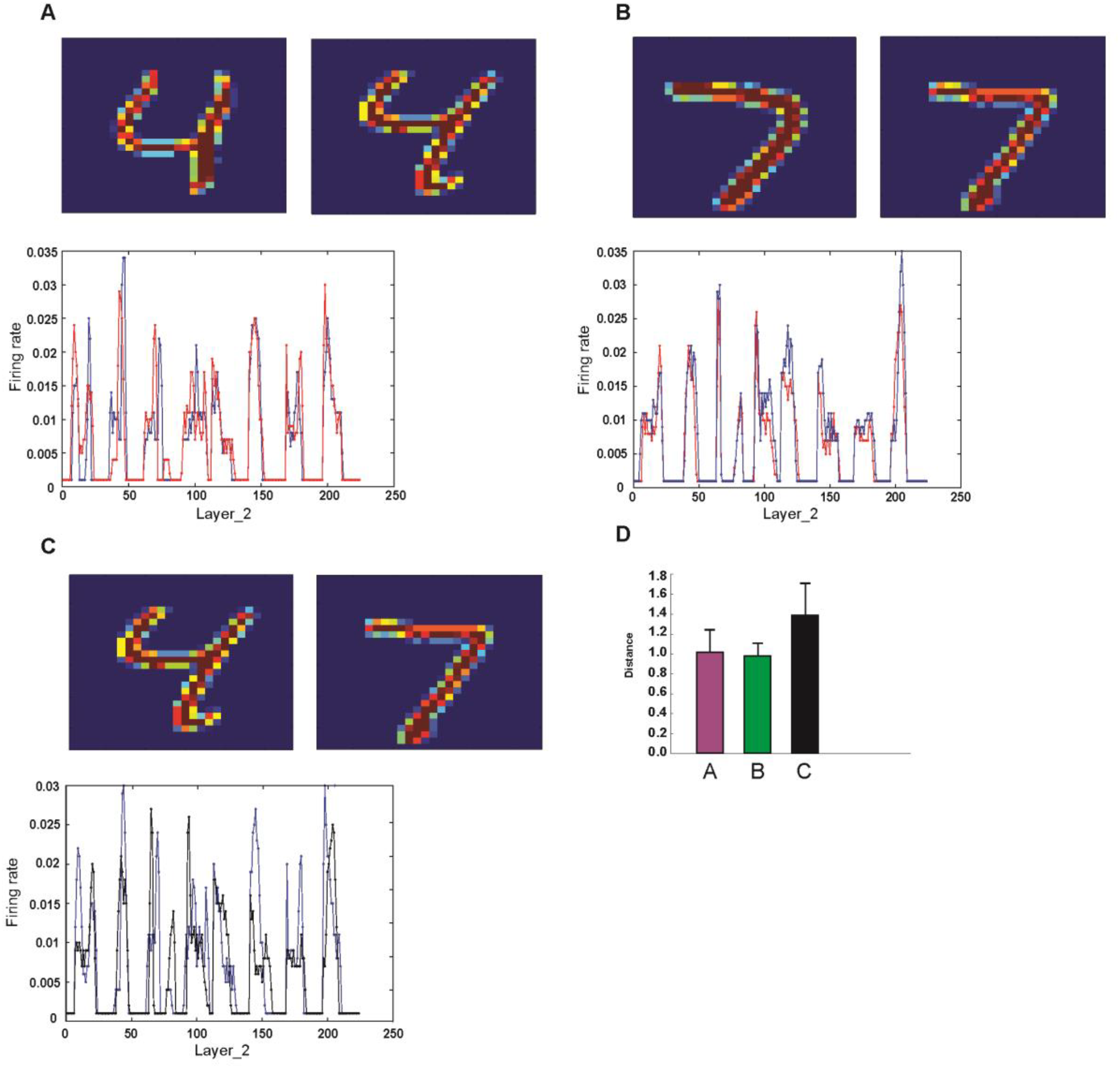
Comparison of neuronal representation of images of different digit classes. **A.** Two samples from the dataset are selected and the induced neuronal activity of the second layer is compared for digit ‘4’ and **B.** for digit ‘7’. **C.** Two samples of digit ‘4’ and ‘7’and their induced neuronal activity of the second layer is compared. **D.** The distance between firing patterns are measured. Images from different digits show higher distance values compared to samples from the same class.

**Figure 4.**
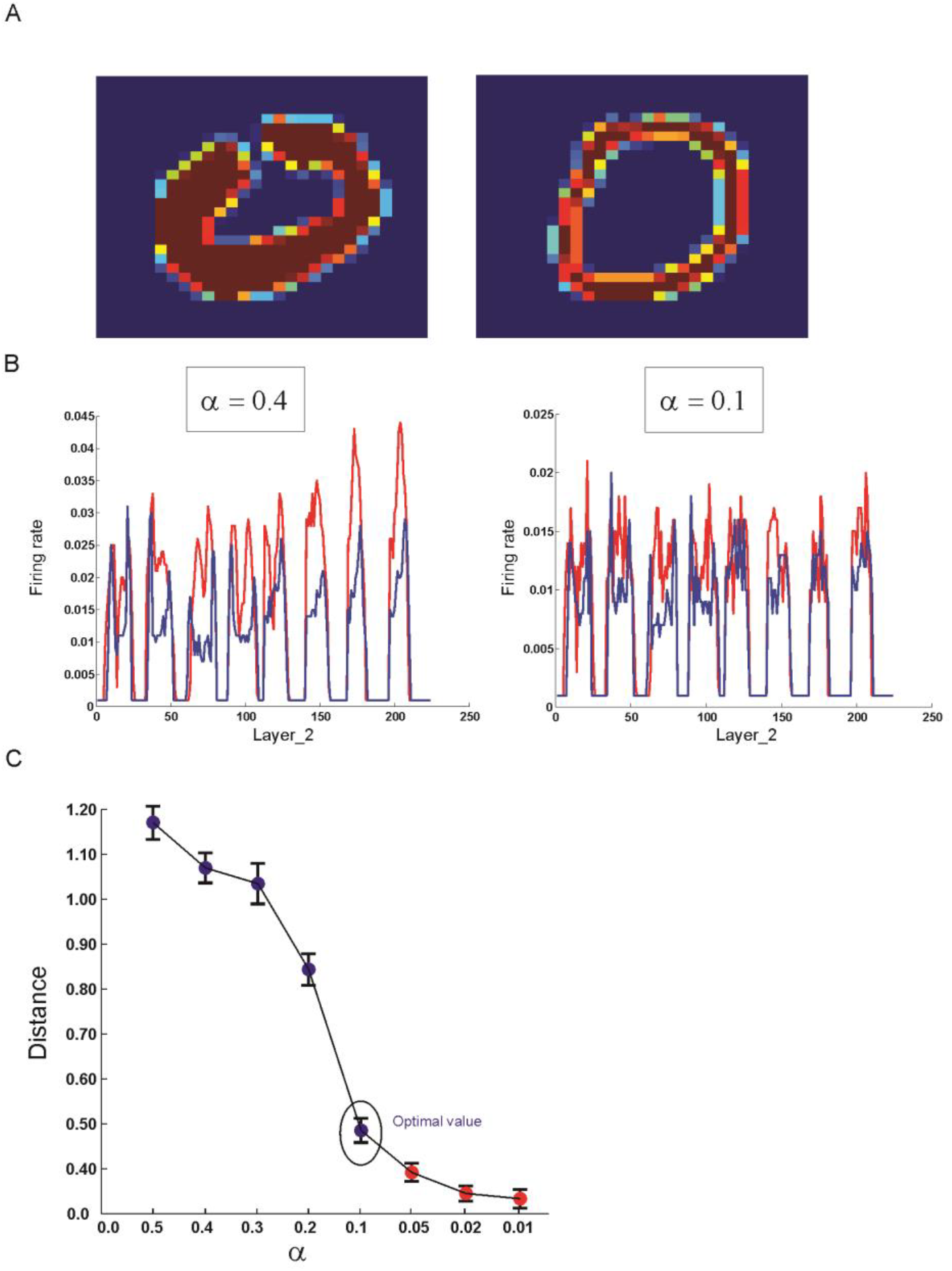
The impact of feed-back inhibition on efficiency of neuronal representation of images. **A.** MNIST database has many samples of hand-written digits with different intensities and this normally makes noise in pattern presentation of images of the same class. In the model, feedback inhibition with different intensities are presented in the second layer such that lower inhibition parameters (α) induce higher inhibition of the second layer (**B**). **C.** Distances between pairs of such images (one thick and one normal written) are measured for different parameter’s value. Optimal parameter value corresponding to minimum average distance without suppressing the neuronal activity of the neurons in the second layer is 0.1.

Figure 4.B demonstrates the activity pattern of neurons in the second layer for two intensity values, lower intensity value, more inhibition on spiking activity of the neurons is applied in the training and test phases. However, increase in the inhibition intensity leads to suppression of spiking activity in the second layer and consequently in the third layer. Therefore, an optimal inhibition intensity were selected as the inhibition value (equal to 0.1) that does not suppress spiking of the third layer and is shown in Figure 4.C. In training phase each training image is presented 12 epochs where at the end of each epoch the synaptic weight between second and third layer is modified according to the learning rule (Figure 5.A). The dynamics of the change in the synaptic weights leads to increase in weights of some neurons that are considered as ‘image features’. At the end of training phase the connections that are less than a threshold (here equal to 0.9) are set to zero (implemented synaptic pruning) (Figure 5.B). For each digit, the training phase is done such that eventually 10 connection matrixes between second and third layers are constructed (Figure 6.A). These connection matrixes named ‘information channels’ in the test phase is used to measure the input into the third layer for each digit to assign one digit to each image. Therefore, one can expect that the similarity between these connection matrixes should be minimized to obtain reliable results. Figure 6.B demonstrate similarity measure for pairs of connection matrixes (left panel) and distance measure for spiking activity of the second layer induced by training images (right panel). The results demonstrates remarkable increase in distance measure by training the network. The performance of the model is calculated by comparing the assigned digits to the images by the SNN and labels of the test dataset. For this purpose, we considered different number of training images for each digit (M=1 to 4) and different pruning thresholds (Figure 7.A). The results show maximum performance (equal to 52%) is assigned to M=2 and optimal pruning threshold equal to 0.8 (Figure 7.B). As the feedback inhibition has shown impact on discrimination of images of the same class with different intensities, the optimal inhibition parameter (***α*** = 0.1) and other values are compared as shown in Figure 7.C. The results show that although feedback inhibition has remarkable impact on discrimination of activity pattern of the second layer induced by images of the same class with different intensities (Figure 4.), however, it has no remarkable impact on the model’s performance. The performance of the model is also studied for each digit individually to show which digit has been recognized efficiently and which digit has misrecognized by the model. Figure 8. demonstrates the model performance for digits 0 to 9 where red bar show percent of the correct recognition while blue bars show the error as misrecognized digits. The results show a very good performance for digits 1,2,4,6,7 and 8 but low performance for digits 0,3,5 and 9. Especially, the model show very low performance to recognize digits 5 and 9. The number of trial for each digit were 2 and the number of epochs were 12.

**Figure 5.**
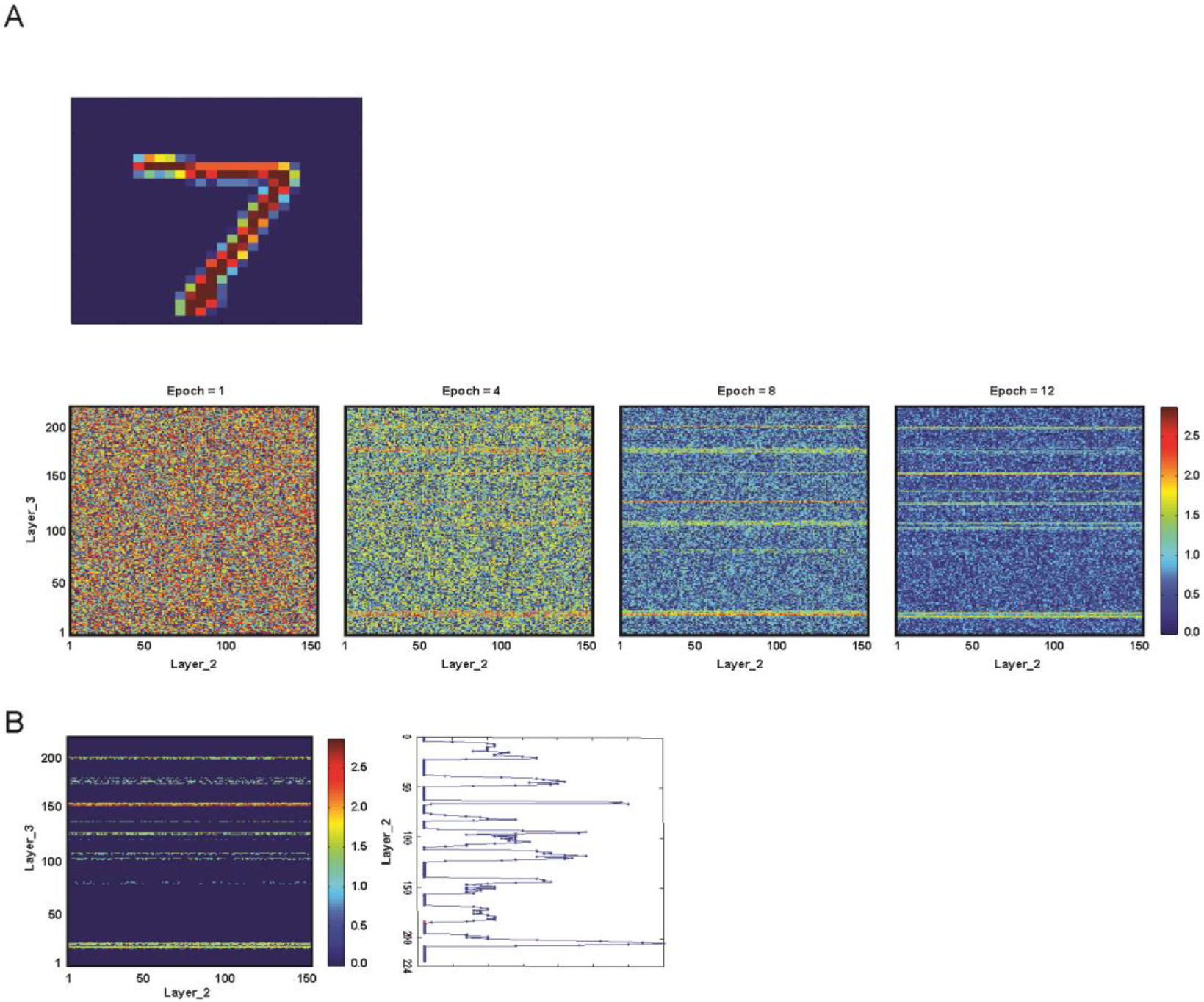
Training of the network with back-propagation and synaptic pruning. **A.** The random synaptic weights between the second layer and third layer is changed according to the back-propagation based learning rule for different epochs of image presentation. An increase in the number of epochs lead to remarkable change in the synaptic weights of highly activated neurons in the second layer and the third neuronal layer. **B.** A pruning threshold (here 0.9) is used to remove weak synapses such that the connection of neurons corresponding to important features of the image is selected.

**Figure 6.**
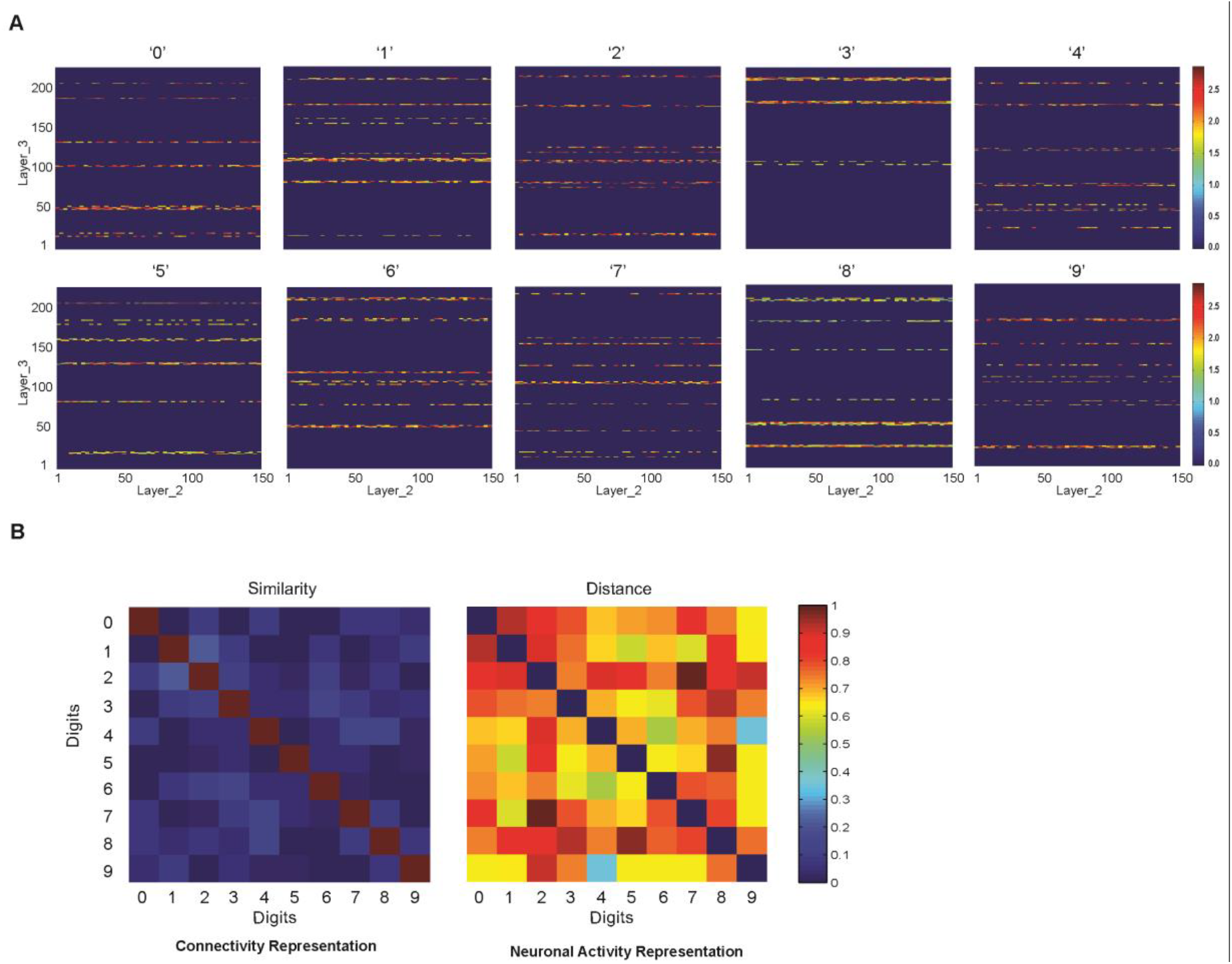
**A.** Connectivity matrixes of digits (0 to 9) after training. Connection and strengths between the second layer and third layer are shown. Number of samples for each digit is one and the epoch equal to 12 were used. **B.** Comparison between neuronal representation of training images (left panel) and similarity of connectivity representation of training images (right panel). The results shows remarkable progress in discrimination of images’ in the network by the proposed training method.

**Figure 7.**
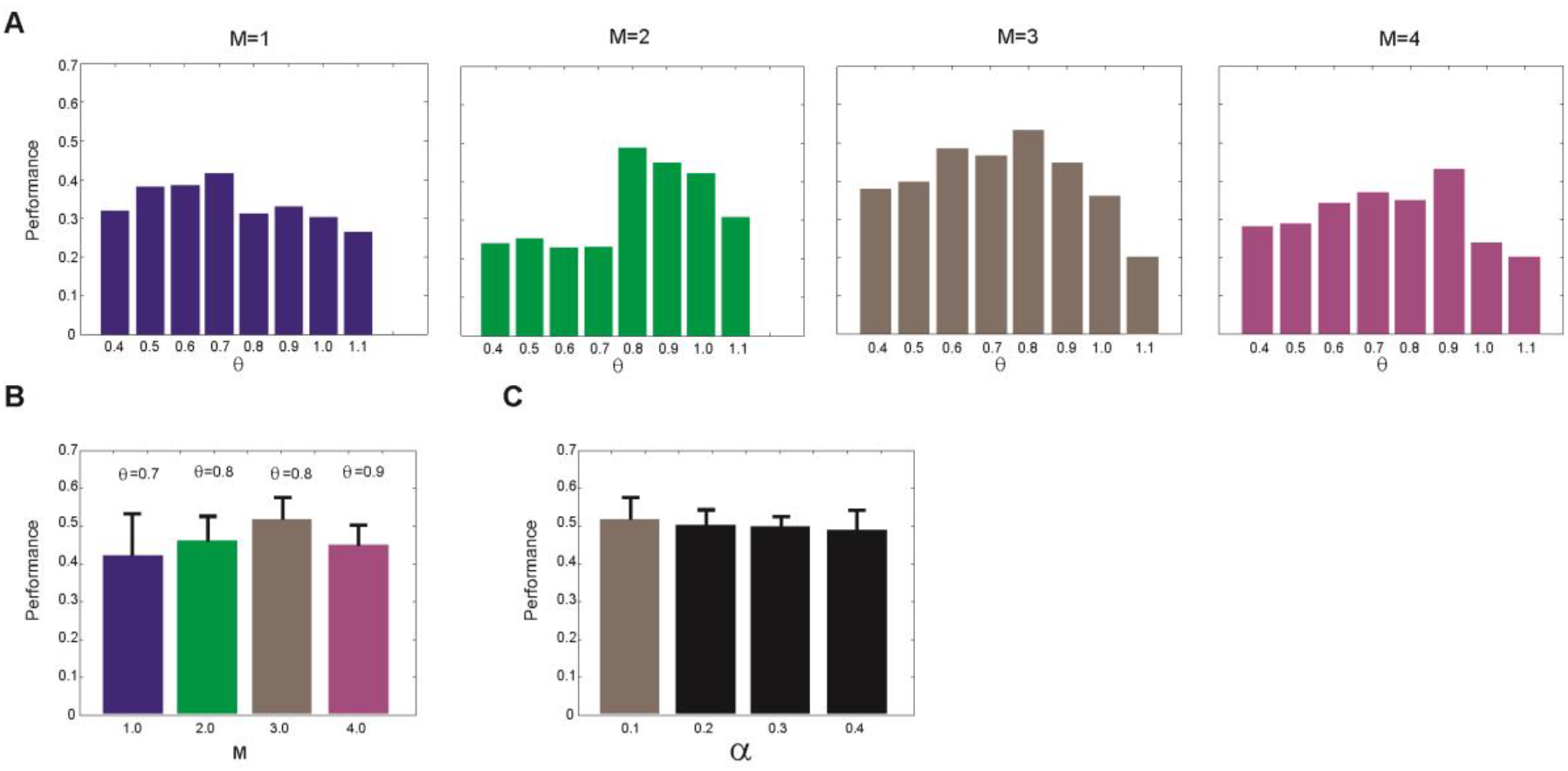
**A.** Classification performance of the method. For the different number of samples (M) for each digit and for different synaptic pruning threshold (θ) the performance is measured. The results shows a shift from synaptic pruning threshold as θ=0.7 (in M=1) to higher values as 0.8 and 0.9 for M=2 to M=4, respectively. **B.** However the maximum performance value for M values is assigned to M=3 at θ=0.8. **C.** The performance of recognition of digits (0 to 9) for different inhibition parameter is shown.

**Figure 8.**
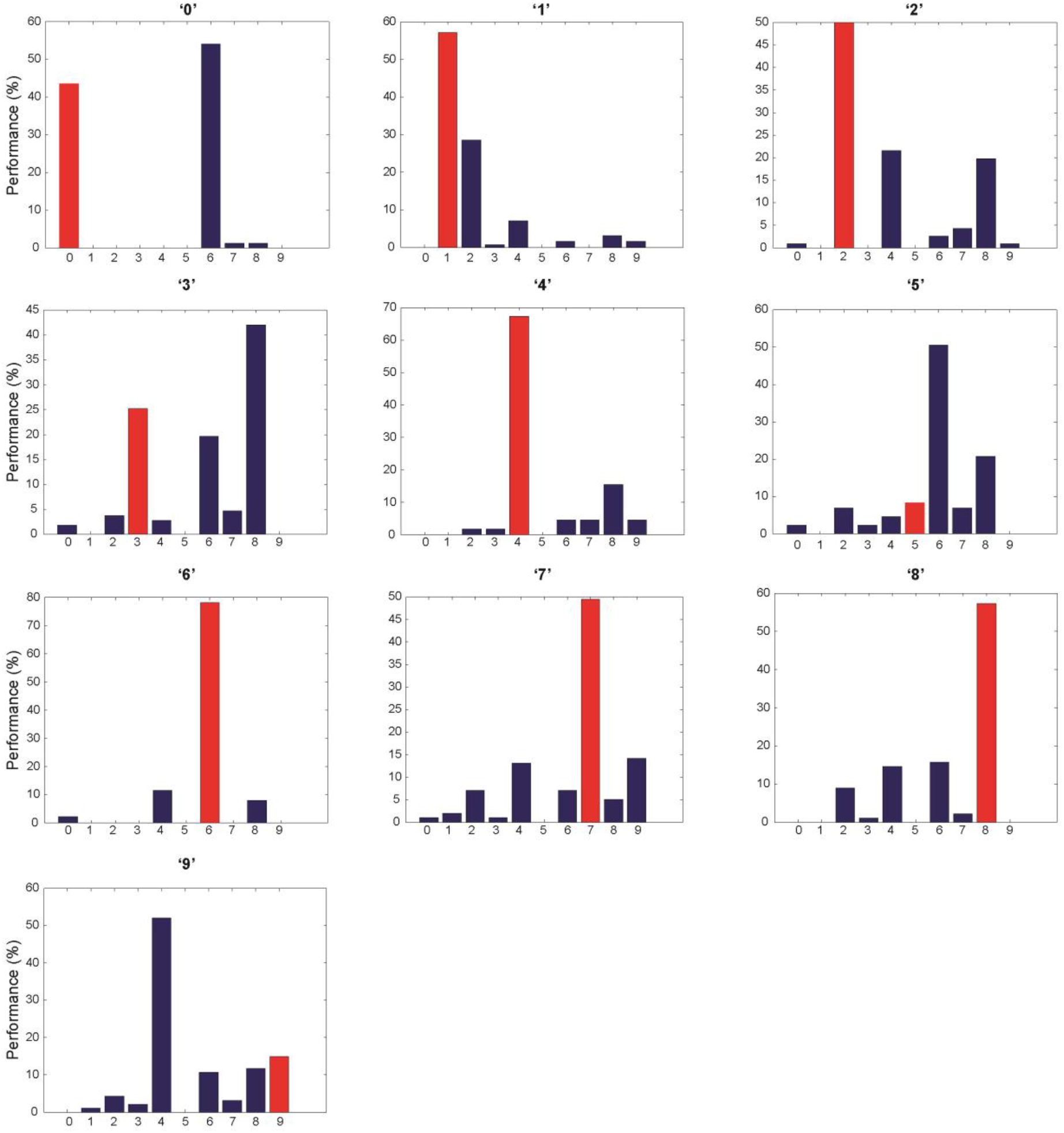
Classification performance for different digits for number of samples for each digit class as M=3 and the threshold of synaptic pruning as θ=0.8. Red bars show the performance for each digit and blue ones are misrecognized digit assigned by the network. Compared to other digits, low performance is obtained for digits 5 and 9.

## Discussion

The capabilities of the human brain to learn and retrieve complex and noisy visual patterns while using small training data have inspired researchers and computer scientists to develop artificial systems that demonstrate similar capabilities. This amazing capabilities originate from neuronal architectures and synaptic mechanisms in different neurons in the brain that are partially known and can be used in developing future artificial systems [32]. Deep learning methods needs large training data in order to show remarkable performance for pattern recognition tasks [33]. However, to apply deep learning models to data analysis, available training data from a few experiments on the observed system is so small that cannot guarantee efficient learning of the network. Therefore, developing deep networks that need training data as small as possible is one of the main challenges of modern machine learning researches. The motivation of our work was to present a hypothesis on how the human brain learn to distinguish hand written digits (as a specific problem of pattern recognition) using a few training samples. For this purpose, we present a novel approach to analysis the digit images to extract features by a few training samples given for each digit.

Modeling of diffusing retrograde signal induced by back-propagation has been used in developing first and second order conditioning in SNNs [34]. In our work, biological back-propagation mechanism and synaptic pruning mechanisms were simplified to present a rate-based learning rule and then implemented in the developed DSNN. To our knowledge, this is the first model that applies inspired biological back-propagation as seen in the neural systems for developing Deep SNNs. Biological back-propagation as a neuronal mechanism play important role in synaptic maturation and consequently normal cognitive functions of the brain [35, 36]. Computer scientists have used the basic mechanism as a synaptic modifying algorithm in artificial neural networks. However, due to discontinuous nature of communication in SNNs, direct implementing of the algorithm in DSNNs is still a challenge but there are some works trying to overcome the challenge by finding learning rules that can achieve high performance as observed in the brains [37–39] or by treating membrane potential of neurons as signals that shows discontinuities at spike times as noise [40].

Synaptic pruning implemented in this work helps the DSNN to be specific to transfer information of images through ‘information channels’ for each digit class (0 to 9) to extract important features representing the images. The implemented synaptic pruning in the model help transform initially fully connected layers to sparse connectivity between second and third layers. Normal synaptic pruning by microglia cells is an essential for neuronal processes in the mouse brain. In addition, contribution of abnormal pruning in neurodevelopmental disorders have been known [41]. For brain-inspired DSNNs, the importance of sparse connectivity in DSNNs for supervised and reinforcement learning has been recently shown while observing high performance on MNIST database [42]. The resulting connectivity matrixes constructed after training named in this work as ‘information channels’ and its existence similarly in biological neural systems and also its potentials for applying in other deep learning architecture (e.g., convolutional neural networks) requires intensive work. Moreover, we expect that this work proposes novel research on developing efficient brain-inspired machine learning methods.

After training, 10 connection matrixes between second and third layers corresponding to digits 0 to 9 are considered as ‘information channels’ to assign a digit to each testing image. Information channels that are constructed by this mechanism through training phase are efficiently being used in recognizing some digits while show low performance for detection of others.

The role of inhibitory neurons and feedback inhibition in efficient pattern separation in hippocampus has been known [43]. Inhibitory neurons have been used in DSNNs to keep excitation-inhibition balance in different deep learning paradigms to speed up training convergence [44, 30].A source of complexity in hand-written digits is the thickness of the images that varies from person to person. An image of a given digit with different size of thickness are presented as different matrixes of values; however, in the visual system these are considered as members of the same class. Such recognition approach is independent of thickness of lines but depends strongly on detection of features. In our work, inhibitory neurons control induced high firing activity of the corresponding neurons in the second layer such that different intensity images (of the same class) are presented by similar matrixes.

In the model, three layers of SNN were used (integrate and fire neuron) and it demonstrate one average 50% performance in digit recognition after training phase. For each digit only 2 training samples and for each sample 12 epochs of training were presented to the network. The performance of our model is about 50%. It means that about half of the testing images were assigned to correct digit that is low comparing to conventional deep learning methods. However, for recognition of digits ‘4’,’6’,’8’ the average performance was about 65%. The results demonstrates the capability of the model to extract horizontal, vertical and orthogonal lines while the model seems to be inefficient in detecting small circles in the images. Therefore, digits ‘1’,’2’,’4’,’6’,’7’ and ‘8’ show performance more or equal to 50%. But digit ‘0’,’3’,’5’ and ‘9’ have not been recognized efficiently. Digit ‘3’ is mainly misrecognized with ‘8’ and digit ‘0’ is misrecognized with ‘6’. Digits ‘5’and ‘9’ are weakly recognized. However, it is important to compare the size of training set (240 training data for each digit class) used in this study and training size of conventional deep learning methods that is up to 15 epoch and each epoch equal to 60,000 samples. The performance of model for different M (number of selected training images to train the network) shows the increase in efficiency by increase in pruning threshold from 0.7 (M=1) to 0.9 (M=3). Any increase in M leads to remarkable decrease in performance. Therefore, these information channels have capacity that is optimal in M=2. In addition, the model is robust to image intensity that we have tried to control its effect on spiking activity of the neurons in the second and third layers.

We think that any better feature extraction method that can overcome challenges to detect lines and circles that compose digits can result in much better performance on digits that are currently being recognized weakly.

In conclusion, we suggest that the basic synaptic and network mechanism used in this DSNN may shed light on how visual system does complicated feature selection efficiently with a few training samples and how to develop new class of artificial systems. This DSNN as a new neuroscience-inspired machine learning method demonstrated efficiency to accomplish complicated tasks where the inputs are subject to noise and uncertainty. We think that in future, models like the SNN developed here may play critical role in developing DSNNs to perform cognitive tasks for modern machine learning techniques.

